# ARID2 loss destabilizes PBAF and drives colorectal cancer

**DOI:** 10.64898/2026.04.01.715786

**Authors:** Sanjana Sarkar, Jimlee Saikia, Murali Dharan Bashyam

## Abstract

The PBAF is one of three biochemically distinct BAF chromatin remodelers in humans. We previously proposed the role of ARID2, a PBAF component, as a bonafide tumor suppressor in colorectal cancer (CRC). Here, we validated loss of tumor suppression under conditions of ARID2 deficiency emanating from a marked reduction in PBAF complex assembly resulting from destabilization of PBAF-specific components BRD7, PHF10, and PBRM1. Transcriptome profiling of ARID2 deficient CRC cells revealed perturbation of disease processes, including CRC and neurodegenerative disorders, as well as CRC relevant pathways including Wnt/β-catenin signalling, but transcript levels of PBAF-specific components remained unchanged, confirmed by RT-qPCR and TCGA data analysis. Our study establishes ARID2 as a critical stabilizer of the PBAF complex of relevance to CRC.

## Introduction

SWI/SNF is a multi-protein complex serving as the primary ATP-dependent chromatin remodeler in eukaryotes (1). It regulates chromatin accessibility to facilitate essential nuclear processes, including transcription (2), DNA repair (2), and RNA splicing (3). The complex utilizes energy derived from ATP hydrolysis, mediated by two mutually exclusive catalytic subunits, BRG1 and BRM, hence its alternative name, the BRG1/BRM-associated factor (BAF) complex, to disrupt histone-DNA interactions (4). The SWI/SNF complex contains three mutually exclusive AT-rich interaction domain (ARID) containing paralogous subunits namely ARID1A, ARID1B, and ARID2. ARID1A/B define the cBAF while ARID2 defines the PBAF complex, respectively (5). ARID2 is an 1,835 amino acid protein with five distinct domains, including the ARID and two additional DNA-binding domains namely a tandem C2H2 zinc-finger domain and an RFX domain in addition to an LXXLL motif and a proline rich domain (6). ARID2 is proposed as a scaffold for the PBAF complex assembly; the SWI/SNF core module interacts with ARID2, followed by the recruitment of BRD7, PHF10, and PBRM1, thereby establishing the PBAF complex (7).

Subunits of the SWI/SNF complex are inactivated by mutations in approximately 25–30% of human cancers, and the complex is widely regarded as a tumor suppressor (8). Unlike other commonly mutated cancer genes, SWI/SNF mutations were discovered relatively recently. Thus, despite their high frequency, the mechanistic links between these mutations and tumorigenesis remain under explored. Our recent exome sequencing analysis of Early-Onset Sporadic Rectal Cancer (EOSRC) identified frequent inactivating mutations in ARID2 (9).

Compared to the BAF (ARID1A/B in particular), the PBAF complex (and ARID2 in particular) is less extensively studied with regard to regulation of nuclear functions. Interest in ARID2 biology increased following its implication in hepatocellular carcinoma (10,11), link to enhancer regulation in neuroblastoma (12) and role in transcriptional repression at DNA double-strand breaks in cooperation with cohesins and PRC2 (13). ARID2 is known to recruit RAD51 for homologous recombination for DNA repair (14).

Here, we demonstrate that loss of ARID2 leads to a marked reduction in the levels of other PBAF components, resulting in improper PBAF complex assembly in CRC cells. Importantly, we show that this reduction happens at the protein (and not at the transcript) level. Furthermore, we identify specific genes and signalling pathways that are differentially regulated upon ARID2 loss and absence of ARID2 appears to drive tumorigenic phenotype in CRC cells grown in culture or as xenografts, providing new insights into the role of ARID2 and the PBAF complex in CRC pathogenesis.

## Materials and methods

### Cell culture and manipulations

The HCT116, SW620, and HT-29 CRC cell lines, procured from the National Center for Cell Science (Pune, India), were authenticated by STR profiling and confirmed to be free of mycoplasma contamination. All cell lines were maintained in Dulbecco’s Modified Eagle’s Medium supplemented with 10% FBS and an antibiotic/antimycotic solution (Thermo Fisher Scientific, Waltham, MA, USA). Transient transfection was carried out on cells at 60–70% confluency using linear polyethylenimine (PEI) at a 1:3 DNA:PEI concentration ratio. Plasmids used for transfection: ARID2-GFP (9); ARID2-SFB and HALO-BAF155 were generated by subcloning the respective cDNAs in the pMH destination vector with the appropriate tags (15).

ARID2 knockout (KO) HCT116 cells were generated using the CRISPR–Cas9 technology as per standard approach (16). Briefly, twenty-base-pair sgRNAs (single-guide RNAs) namely GTAAGCCAGCCAGCTCAACA (targeting exon 15; Broad Institute: https://portals.broadinstitute.org/gpp/public/analysis-tools/sgrna-design), and GCTATACATGCTCACGGAAATGG (targeting exon 10; (17)) to guide specific Cas9 mediated cleavage. The sgRNAs were cloned into a pU6-Cas9-GFP vector (a kind gift from Dr. P. Chandra Shekar, CSIR-Centre for Cellular and Molecular Biology, Hyderabad, India). The resulting recombinant plasmids were transfected into HCT116 cells and GFP-positive cells were FACS-sorted and serially diluted for single-clone selection. The KO was confirmed by both DNA sequencing and immunoblotting. We selected two clones KO1 (targeting exon 10) and KO5 (targeting exon 15).

Cell-line manipulations including cell growth, viability (3-[4,5-dimethylthiazol-2-yl]-2,5 diphenyl tetrazolium bromide (MTT)) and colony formation, were performed as described previously (18). RT-qPCR (primer details in Table S1), immunoblotting, and protein pull-down and immunoblotting assays were performed as described earlier (19). For proteasome inhibition, cells were treated with MG132 (Cat no. C2211, Sigma-Aldrich, St. Louis, MO, USA) (10 μM) for 6 h and subjected to immunoblotting. For localization studies, immunofluorescence was carried out as previously described (19), and the slides were visualized at 65× magnification using an LSM 900 confocal microscope (Carl Zeiss AG, Oberkochen, Germany).

### Nude mice xenograft experiments

Following approval from the BRIC-CDFD Institutional Animal Ethics Committee (protocol EAF/MDB/12/2025), 7-8-week-old female Foxn1−/− nude mice (n = 6) were randomly selected and subcutaneously injected with HCT116 or HCT116 ARID2 KO cells (2×10^6^ cells each). Tumor diameters were measured weekly from week 3 using a Vernier caliper, and volumes were calculated as (W^2^ × L)/2. Mice were euthanized 5 weeks post-injection, and tumors were excised for measurement of weight and volume.

### RNA-seq

#### RNA Extraction and Library Preparation

Total RNA was extracted from CRC cell lines using the TRIzol reagent (Life Technologies, Grand Island, NY, USA) followed by purification with the RNeasy Kit (Qiagen, Valencia, CA, USA) according to the manufacturer’s protocols. Transcriptome libraries were constructed using the NEBNext Ultra II Directional RNA Library Prep Kit (New England Biolabs, Ipswich, MA, USA) with poly(A) mRNA enrichment prior to library preparation.

#### RNA Sequencing

For knockout (KO), sequencing was performed on the NovaSeq X Plus platform (Illumina, San Diego, CA, USA) using paired-end sequencing with a read length of 100 bp (n = 3 biological replicates). For knockdown (KD), sequencing was performed on the NextSeq 2000 platform (Illumina) using similar approach but single-end chemistry (n = 2 biological replicates).

#### RNA-seq Data Processing and Alignment

Raw read quality was assessed using FastQC (20). For KD samples, reads were aligned to the human reference genome (GRCh38, Ensembl release 115) using STAR aligner, and gene-level read counts were obtained using featureCounts as described previously (21). For KO samples, adapter trimming was performed using Cutadapt (22) prior to quantification. Trimmed reads were pseudo-aligned and quantified at the transcript level using Salmon (23) against the GRCh38 reference transcriptome (Ensembl release 115), and transcript-level abundances were aggregated to gene-level counts using tximport (24).

#### Differential Gene Expression Analysis

Differential gene expression (DGE) analysis was performed using DESeq2 (25) in R. Genes were considered significantly differentially expressed at an adjusted p-value < 0.01 (Benjamini–Hochberg FDR < 10%) and an absolute log_2_ fold change ≥ 0.5.

#### Functional Enrichment Analysis

Gene Ontology (GO) term and KEGG pathway enrichment analyses were performed in R using the clusterProfiler package (enrichGO and enrichKEGG functions; (26)) on significantly upregulated and downregulated genes (identified using DESeq2), separately. Enrichment analysis was also performed for the upregulated genes for OMIM_Disease against DAVID annotation datasets (27) using default parameters. Gene Set Enrichment Analysis (GSEA) was conducted using GSEA software (v4.1.0;(28)) with the MSigDB Hallmark gene set collection as reference datasets. Top enriched pathways were selected based on normalized enrichment score (NES) and FDR q-value < 0.25 as per standard GSEA thresholds.

### Antibodies

Primary antibodies used for pulldown, immunoblotting and immunofluorescence (IF) were ARID2 (1:5000, Cat no. 82342, Cell Signaling Technology, Danvers, MA, USA), BRD7 (1:5000, Cat no. A302-304A, Bethyl Laboratories, Montgomery, TX, USA), PBRM1 (1:5000, Cat no. A301-591A, Bethyl Laboratories), PHF10 (1:5000, Cat no. PA5-30678, Invitrogen, Carlsbad, CA, USA), BAF155 (1:5000, Cat no. sc-32763, Santa Cruz Biotechnology, Dallas, TX, USA), BRG1 (1:5000, Cat no. sc-17796, Santa Cruz Biotechnology), SMARCD1 (1:5000, Cat no. HPA004101, Sigma-Aldrich), ARID1A (1:5000, Cat no. A19570, ABclonal Technology, Woburn, MA, USA), SS18 (1:5000, Cat no. A6990, ABclonal Technology), Lamin B1(1:5000, Cat no. 12586, Cell Signalling Technology), GAPDH (1:10000, Cat no.MAB932Hu22, Cloud-Clone Corp, Katy, TX, USA), Anti-Halo (1:10000; Cat no. G9211; Lot no. 0000348664; Promega, Madison, WI, USA), Anti-Flag (1: 10000; Cat no. F1804; Lot No. SLBW5142; Sigma-Aldrich).

Secondary antibodies used were horseradish peroxidase (HRP)-conjugated goat anti-rabbit (1:10,000; Cat no. 65-6120, Invitrogen), HRP-conjugated goat anti-mouse (1: 10,000; Cat no. 62-6520, Invitrogen), and Alexa Fluor 568-conjugated anti-rabbit (1:400; Cat no. A11036, Thermo Fisher Scientific).

## Results

### ARID2 loss drives colorectal carcinogenesis

To validate ARID2’s impact on CRC tumorigenesis, we generated ARID2 KO CRC cells (Fig 1A). RT-qPCR for canonical tumor suppressive *ARID2* target genes, namely *BMP4* and *CDKN1B* (Fig 1B), and tumorigenic assays including cell viability (MTT), growth, colony formation (Fig 1C) and nude mice xenografts (Fig 1D), revealed ARID2’s possible role as a tumor suppressor in CRC, consistent with our earlier results (9). Further, reintroducing ARID2 in ARID2 deficient background restored the phenotype observed in wild type cells, confirmed by multiple tumorigenic assays (Fig. 1E), further validating ARID2’s role in tumor suppression.

**Figure 1.**
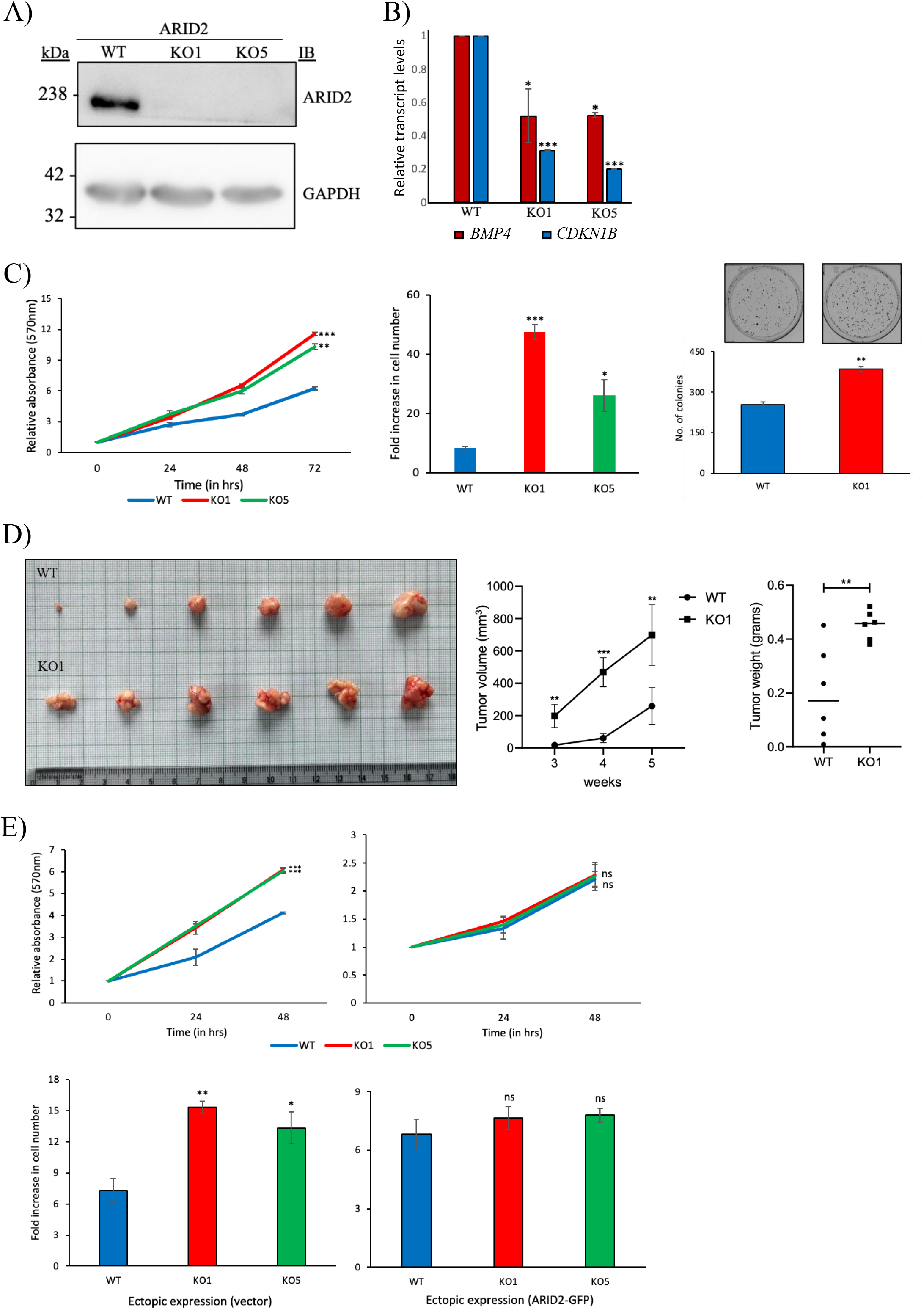
ARID2 loss drives colorectal carcinogenesis: A) Validation of ARID2 depletion in HCT116 ARID2 knockout cells using immunoblotting. GAPDH was utilized as loading control. B) RT-qPCR analysis demonstrates significantly reduced mRNA expression of classical PBAF-regulated target genes *BMP4* and *CDKN1B* in ARID2 KO cells. C) Tumorigenic potential of ARID2 proficient and deficient HCT116 cells was assessed via cell viability (left), growth (middle; measured after 72 hours), colony formation (right) assays. D) ARID2 deficiency accelerates tumor growth as determined by nude mice xenograft assays. Representative images of excised xenograft tumors harvested five weeks post-injection (left) and quantitation of volume (middle) and weight (right) are shown. WT: wild type ARID2; KO (KO1): ARID2 knock out. E) Ectopic expression of ARID2 rescues tumor suppressor function in ARID2 KO cells. Tumorigenic potential was assessed via cell viability (top) and growth (bottom) assays. All results are from three independent experiments and are presented as mean ± SD. Statistical significance was determined using unpaired Student’s t-test; *P<0.05; **P<0.01; ***P<0.001; ns, not significant.

### Loss of ARID2 disrupts PBAF complex integrity at protein level

ARID2 is known to support PBAF complex assembly (7). We therefore evaluated PBAF complex formation based on affinity-pull down of a cBAF/PBAF/ncBAF core component namely Halo-tagged BAF155, in ARID2 proficient and deficient CRC cells. PBAF complexes could not be isolated in the ARID2-KO cells, though the same could be readily isolated in ARID2 proficient cells (Fig 2A). Of note, no perturbation of cBAF complex formation was revealed (Fig 2A).

**Figure 2.**
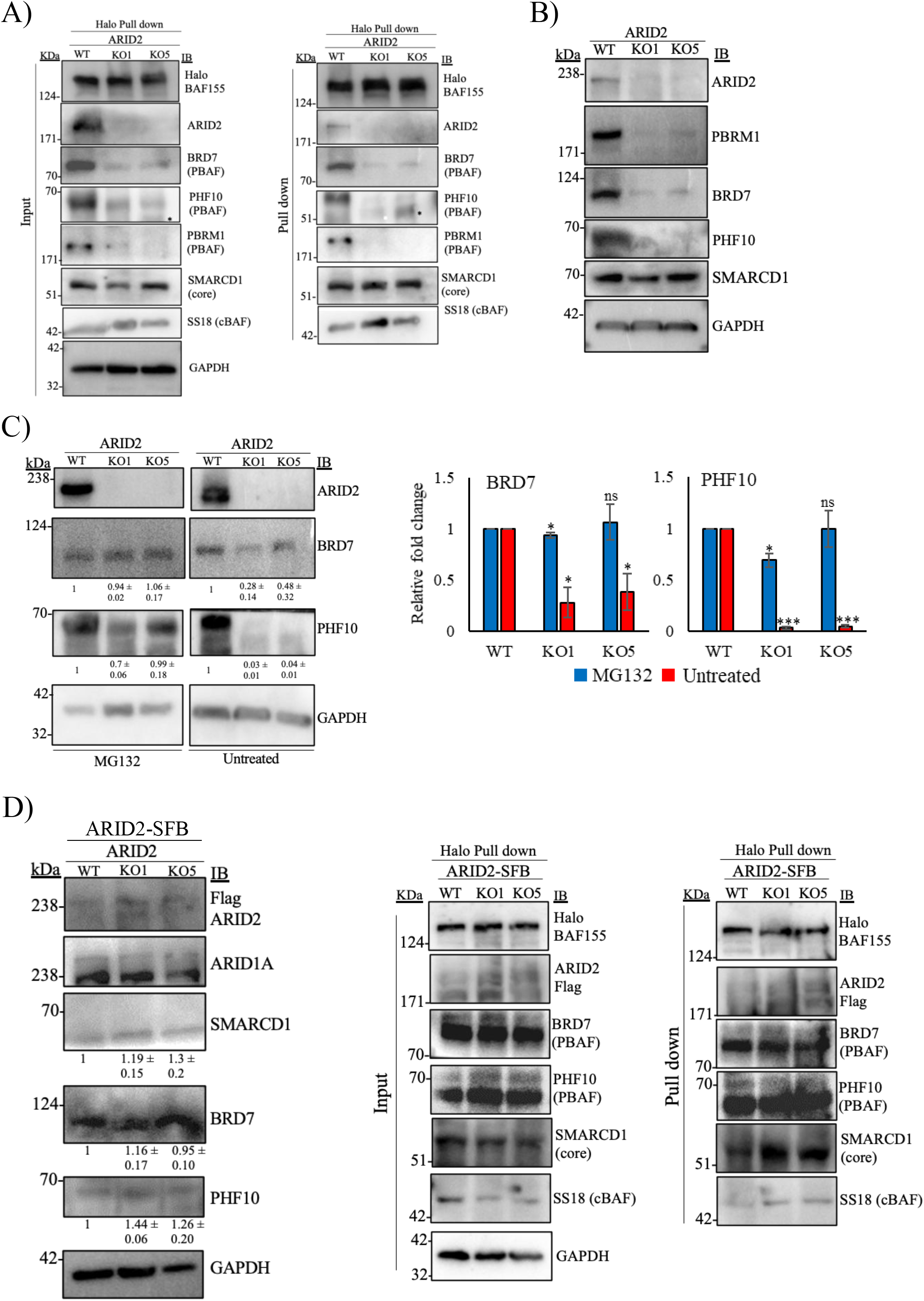
Loss of ARID2 disrupts PBAF complex integrity at protein level: A) Evaluation of PBAF complex formation ability of ARID2 proficient and deficient HCT116 cells based on affinity purification of Halo-tagged BAF155 followed by immunoblotting. *; non-specific band. B) Immunoblot analysis of SWI/SNF subunit expression in wild type and ARID2 deficient (KO1, KO5) HCT116 cells. GAPDH was utilized as loading control. C) Immunoblot and corresponding densitometric quantification of PBAF components BRD7 and PHF10 levels in ARID2 proficient and deficient HCT116 cells treated with the proteasome inhibitor MG132 (10µM). GAPDH was utilized as loading control. Results are quantified from three independent experiments and presented as mean ± SD. D) Ectopic expression of ARID2-SFB in ARID2 deficient (KO) HCT116 cells successfully rescues the protein levels of endogenous PBAF subunits BRD7 and PHF10 (left). GAPDH was utilized as loading control. ARID2-SFB rescues PBAF complex formation in ARID2 deficient (KO) HCT116 cells as determined by affinity purification of Halo-tagged BAF155 (right).

The inability to detect PBAF complexes in ARID2-KO CRC cells could stem from a possible destabilization of PBAF protein components. Indeed, ARID2 KO cells exhibited reduced levels of PBAF-specific components (BRD7, PHF10, and PBRM1) (Fig. 2B), while core component (SMARCD1) remained unchanged (Fig 2B). Further, we observed similar results in CRC cell lines (HT-29 and SW620) exhibiting ARID2 KD (Fig. S1A). These observations were confirmed by immunofluorescence assays (Fig S1 B). In order to determine the basis for reduction of PBAF component levels, the ARID2 deficient cells were treated with MG132, a proteasomal inhibitor. MG132 treatment, rescued the protein levels of PBAF components in ARID2 deficient cells (Fig 2C). Finally, ectopic expression of ARID2 in ARID2 deficient cells restored BRD7, PHF10, and PBRM1 levels (Fig. 2D) besides reinstating PBAF complex forming ability.

### Perturbation of tumorigenesis pathways upon ARID2 loss in CRC cells

We performed transcriptome profiling of ARID2-KO HCT116 CRC cells and of HT-29 and SW620 CRC cells (Fig S2A) exhibiting ARID2 KD (9). Results revealed a) robust suppression of cell processes such as cell cycle check points, chromosomal segregation, and DNA damage response (Fig. 3A), b) upregulation of disease phenotypes (Fig 3A) including colorectal and other cancers (Fig 3B), and dysregulation of signalling pathways including NFKB, TNF-beta, and Wnt/β-catenin (Fig S2B and Fig S2C), under conditions of ARID2 deficiency. Of note, Wnt/β-catenin signalling is a prominent canonical driver of CRC initiation and progression (29). Multiple cancer-related genes were differentially expressed including E2F targets (*MYC, MCM6, MCM7*, and *CCNE1*), epithelial mesenchymal transition (EMT) genes (*VIM, TGFB1*, and *ACTA2*) and Wnt/β-catenin signalling genes (*DKK4, WNT5B, CCND2*, and *MYC)* (Fig S2D). Finally, analysis of the RNA Seq data revealed downregulation of the classical tumor suppressive *ARID2* target gene *CDKN1B* (Fig 3C) in ARID2 KO CRC cells. Both *CDKN1B* and another canonical tumor suppressive *ARID2* target namely *BMP4* were confirmed to be downregulated, in ARID2 deficient (both KO and KD) cells, by RT-qPCR (Fig 3D) (9). These results underscore the functional role of ARID2 as a critical tumor suppressor in colorectal malignancies.

**Figure 3.**
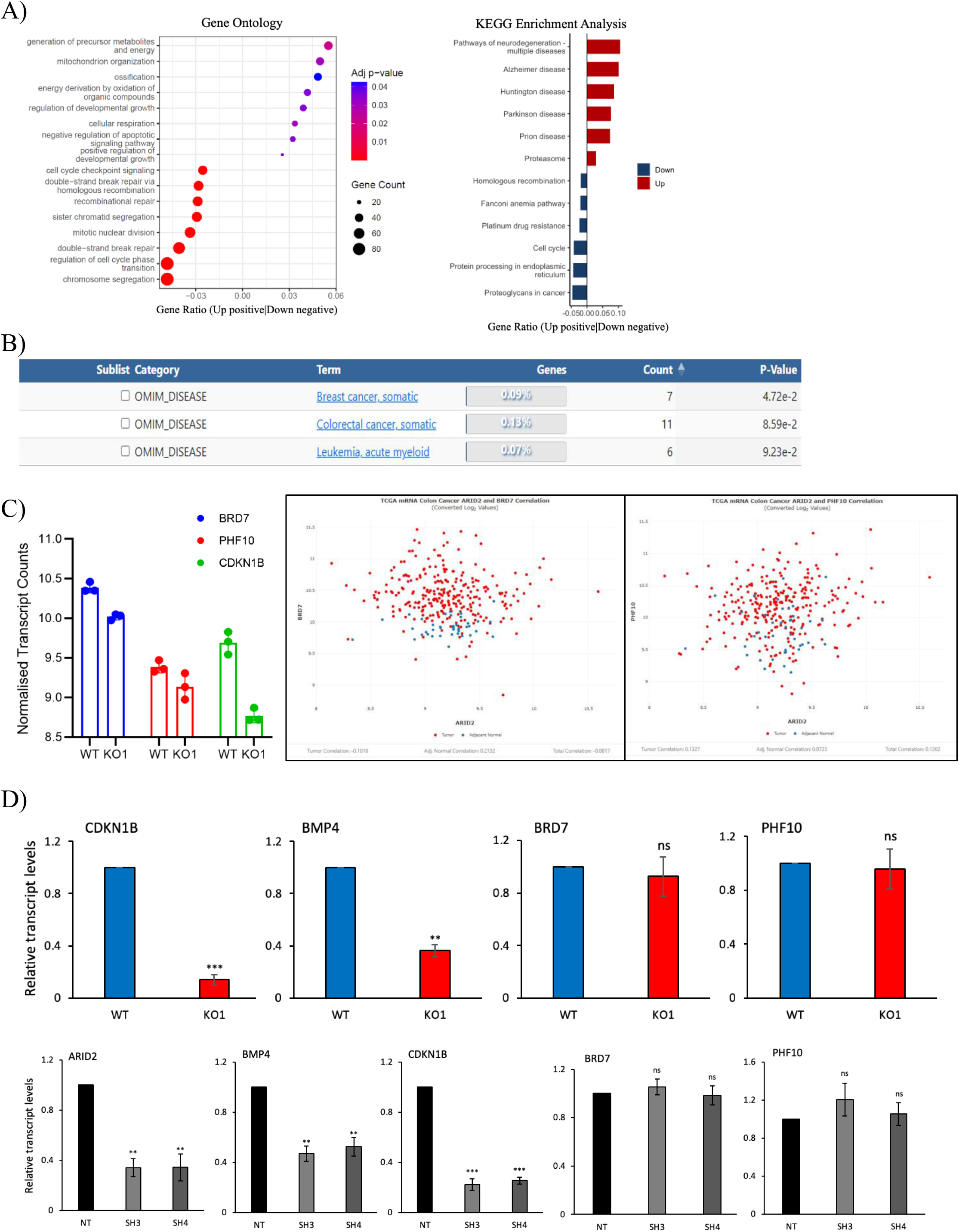
Identification of deregulated gene expression profile upon AIRD2 loss in CRC: A) Functional enrichment analysis of the ARID2-dependent transcriptome in HCT116 cells. Gene Ontology (GO) for biological processes (left) and KEGG pathway (right) analyses of differentially expressed genes in ARID2 deficient (KO1) vs. proficient (wild type) HCT116 cells is shown. Statistical thresholds for selected gene sets were set at an adjusted p-value < 0.01, False Discovery Rate (FDR) < 10%, and ± 0.5 log_2_fold change in knockout with respect to WT. B) Enrichment analysis for differentially upregulated genes using the OMIM_Disease panel in the DAVID annotation datasets using default settings. C) Expression correlation between PBAF subunit transcript levels in RNA seq performed on ARID2 KO vs wild type HCT116 cells (left; bar plots represent normalised transcript counts) and from the TCGA CRC expression data set (right; scatter plots represent mRNA expression correlations). TCGA data were retrieved and cross-referenced via the Proteomics Data Commons (PDC) (https://proteomic.datacommons.cancer.gov/pdc/). D) Quantitative validation of PBAF and ARID2-regulated canonical transcripts in ARID2 proficient and deficient HCT-116 KO cells (top) and in SW620 KD cells (bottom). All results are from three independent experiments and are presented as mean ± SD. Statistical significance was determined using unpaired Student’s t-test; *, P<0.05; **P<0.01; ***P<0.001; ns, not significant.

Interestingly, analysis of RNA seq data of ARID2 KO CRC cells revealed no significant change in transcript levels of *BRD7* and *PHF10* (Fig 3D), validated using RT-qPCR analysis in ARID2 KO and KD CRC cells (Fig 3D). Further, there was no significant correlation of *ARID2* transcript levels with that of *BRD7* and *PHF10* in The Cancer Genome Atlas (TCGA) CRC RNA seq datasets (Fig 3C). These results revealed that loss of ARID2 perturbs PBAF components at the level of protein stability and not at the level of transcription.

## Discussion and Conclusion

*ARID2* inactivation appears to be a frequent event in oral (30), hepatocellular (10), and lung cancer (31). In microsatellite-unstable colorectal cancer (CRC), genes encoding ARID family proteins exhibited high mutation rates of 39% for ARID1A and 13% for ARID1B and ARID2, respectively (32). We previously identified *ARID2* as a tumor suppressor in EOSRC (9). Here, we show that *ARID2* loss results in reduced expression of known tumor suppressive target genes (Fig 1B) and increases tumorigenic potential of CRC cells (Fig 1C). Consistently, complete *ARID2* loss was observed in 18% of CRC TMA samples (9).

ARID2 is known to be the initial subunit that associates with the core SWI/SNF components, thereby driving assembly of the PBAF complex (7). In melanoma, loss of ARID2 leads to reduced levels of other PBAF components and the PBAF complex is also destabilised (33). This study is the first to demonstrate that, in the absence of ARID2, other PBAF subunits become unstable, leading to impaired PBAF complex assembly, contributing to CRC (Fig 2A). The loss of ARID2 is expected to have additional effects (perturbed DNA repair, for example (13), predicted to drive tumorigenesis, not evaluated in this study. Of note, transcriptome analysis of cells exhibiting ARID2 deficiency revealed down regulation of several DNA damage response pathways (Fig 3A), commensurate with ARID2’s known role in regulating DNA repair (14). The analysis also revealed upregulation of multiple neurodegenerative disorders and other related diseases such as Alzheimers, Huntington, and Parkinsons, (Fig 3A); previous studies have validated the role of ARID2 in these diseases (34,35).

We further show that downregulation of PBAF subunits under conditions of ARID2 deficiency occurs at the protein level with negligible contribution at the transcript level. While SWI/SNF subunits are well known regulators of global gene expression, evidence for direct transcriptional cross regulation within the complex components themselves remains sparse. Only one study revealed a possible correlation between wild-type BRG1 overexpression and increased BRM transcripts (36).

In summary, our data explores the mechanistic basis of *ARID2*-loss mediated tumor promotion in CRC (Fig 4). ARID2 deficiency leads to reduction of PBAF component levels. It will be interesting to evaluate a possible therapeutic potential of PBAF destabilization, given that a significant proportion of CRC patient samples exhibit ARID2 inactivation (18%; (9)). For example, it is possible that critical nuclear function(s) are taken over by alternative complexes upon PBAF inactivation, providing avenues to target the compensating complex (37). Targeting synthetic lethality associated with ARID2-deficient CRC, such as inhibition (Tazemetostat) of EZH2 (38) or of ATR/PARP (17), is already being evaluated as therapeutic options exploiting the unique DNA repair and epigenetic vulnerabilities created by PBAF loss in other cancer types (33). Further investigation into ARID2/PBAF associated pathways may reveal additional therapeutic vulnerabilities in CRC.

**Figure 4.**
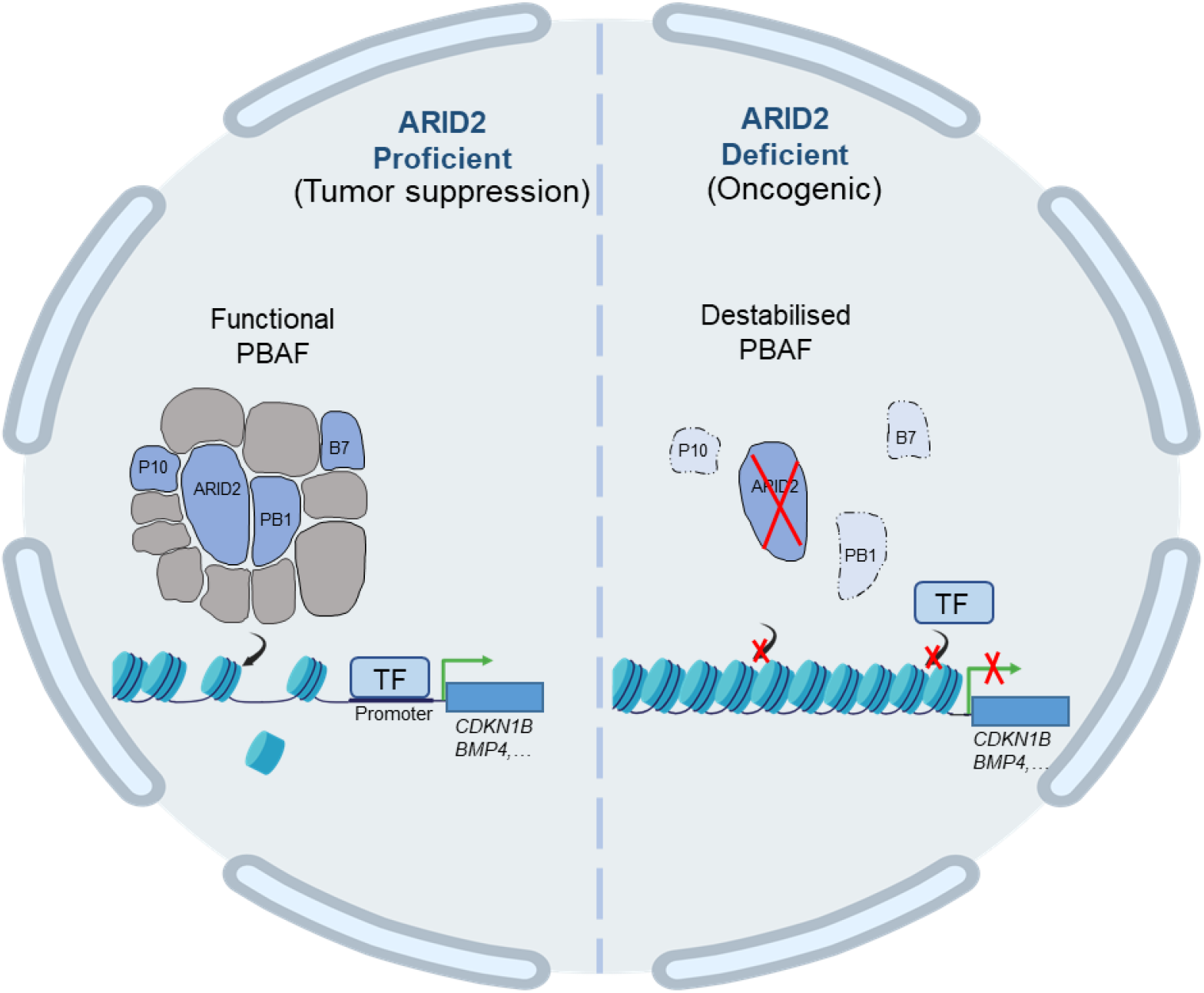
ARID2 deficiency destabilizes PBAF complex leading to CRC progression. The nuclear function of ARID2 is to drive functional PBAF complex formation thereby inducing transcription of classical tumor suppressor genes. In contrast, in the absence of ARID2, the PBAF complex assembly is hampered, negatively impacting the transcription of these genes leading to carcinogenesis in CRC. TF, transcription factor; B7, BRD7; P10, PHF10; PB1, PBRM1. PBAF is adapted from (39). Figure created using BioRender (https://app.biorender.com/illustrations/canvas-beta/699f06ef7b2f44ebc964ba83).

## Author Contributions

**SS**: Conceptualization, Methodology, Formal analysis, Validation, Data Curation, Visualization, Writing-Original draft, Writing - Review & Editing. **JS**: Methodology, Validation, Visualization. **MDB**: Conceptualization, Resources, Formal analysis, Visualization, Supervision, Project administration, Funding acquisition, Writing - Review & Editing.

## Acknowledgements

This research was supported by the DBT/Welcome Trust - India Alliance -Team Science Grant (IA/TSG/23/1/600489) and BRIC-CDFD core funding to MDB. SS, a PhD student at Manipal Academy of Higher Education, acknowledges the Council of Scientific & Industrial Research for junior and senior research fellowships. We thank Dr Apuratha Pandiyan and Ms. Sumaiya Sabnam, Laboratory of Molecular Oncology, BRIC-CDFD, Hyderabad, India, for differential gene expression analyses of RNA Seq data. Additionally, we thank BRIC-CDFD’s Sophisticated Equipment Facility for fluorescence microscopy and Sanger sequencing, and the Experimental Animal Facility for nude mice xenograft experiments.

## Supplementary Figures

**Figure S1.**
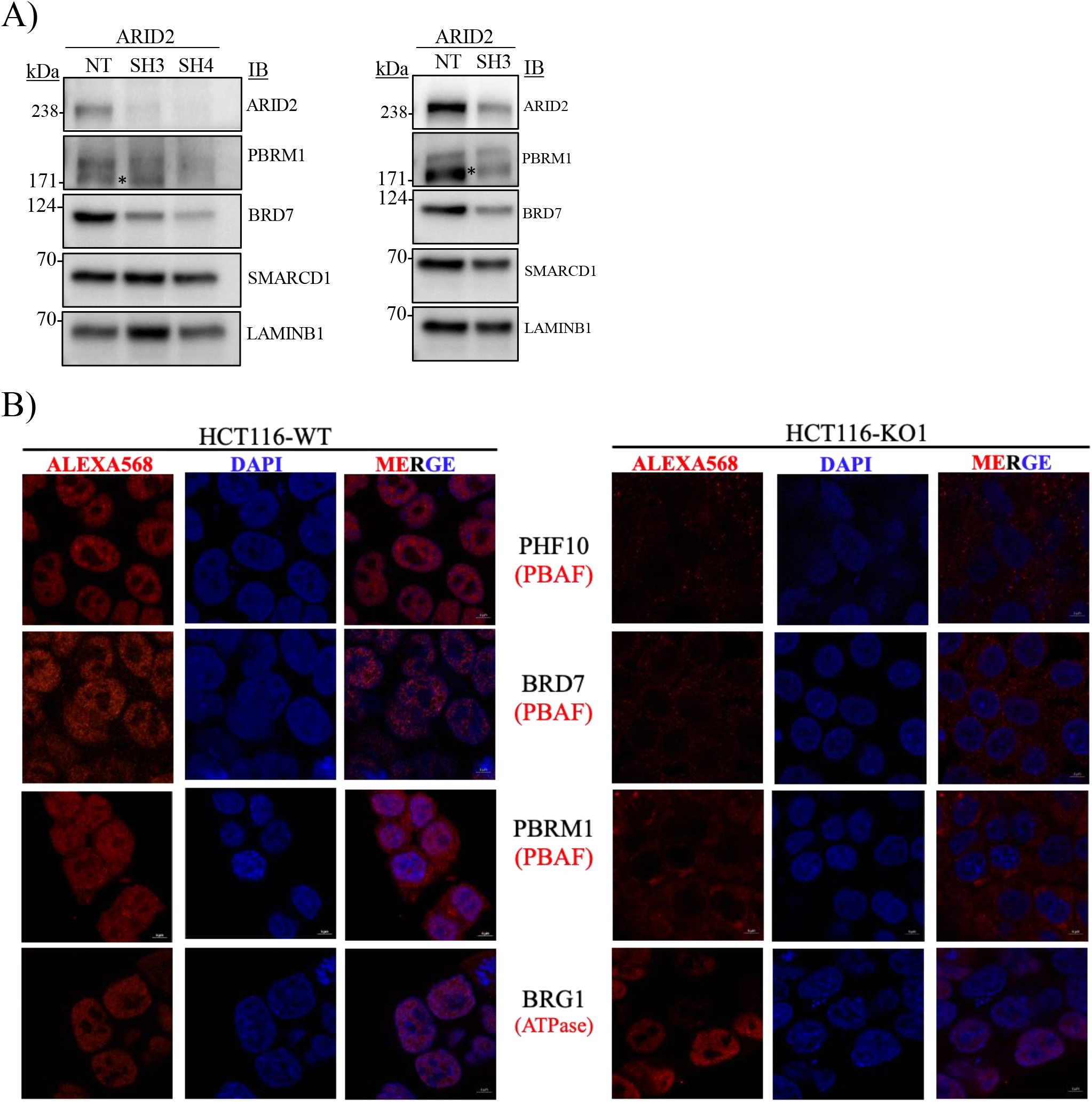
(in relation to Figure 2): A) Immunoblot analysis of SWI/SNF core and PBAF components in HT-29 and SW620 cells following shRNA-mediated ARID2 knockdown (SH3, SH4) compared to non-targeting (NT) controls. Lamin B1 was utilized as loading control. *, non-specific band. B) Visualization of PBAF subunit levels determined via immunofluorescence in the same cells as Panel A. Cells were probed for endogenous PBAF and Pan-BAF (ATPase) components (Alexa Fluor 568, red) and counterstained with DAPI (blue) to stain nuclei. Scale bar 5µm.

**Figure S2.**
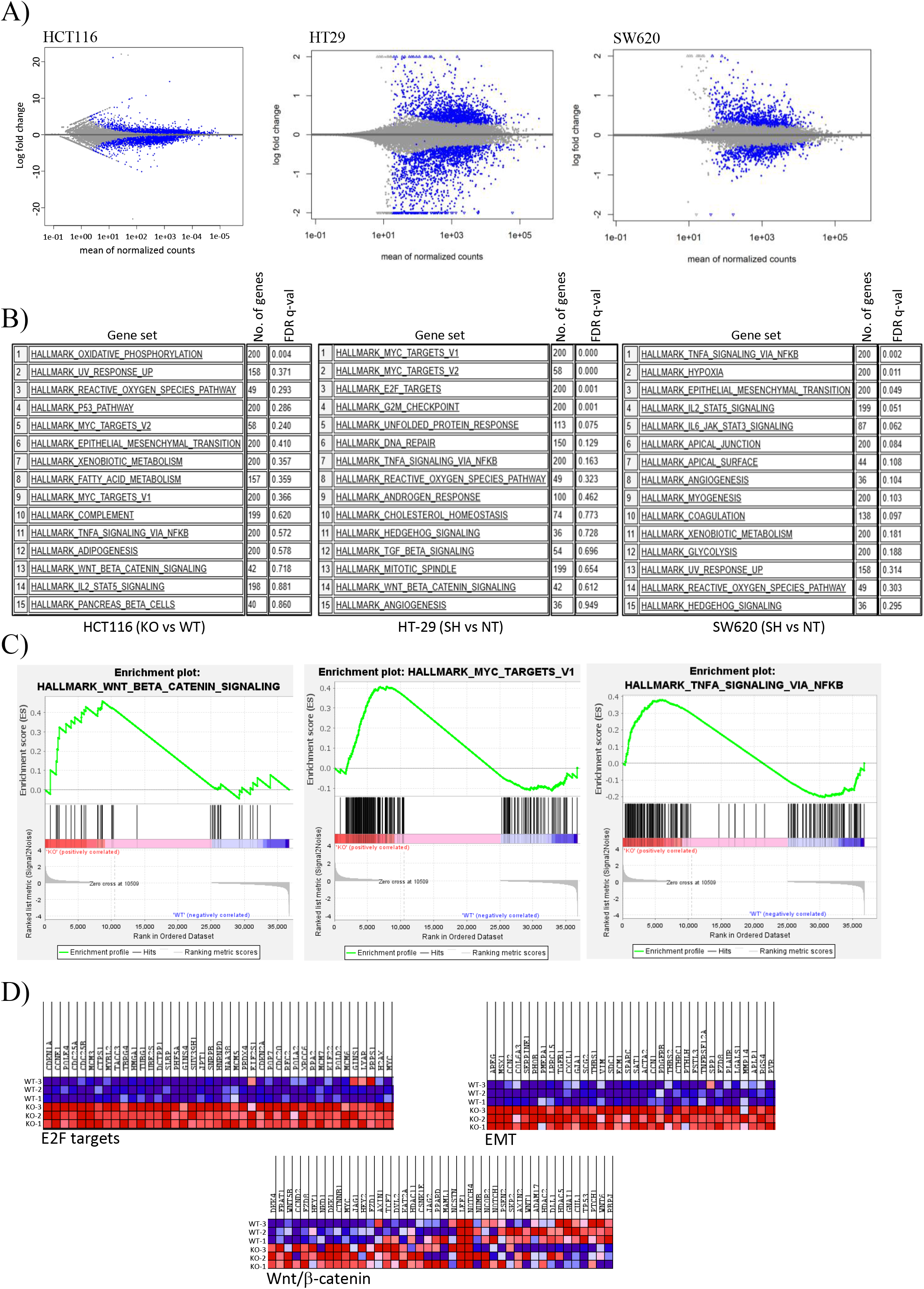
(in relation to Figure 3): A) MA-plots illustrate the distribution of differentially expressed genes in HCT116, HT-29, and SW620 cell lines following ARID2 depletion. Blue data points indicate significantly dysregulated transcripts. B) Gene Set Enrichment Analysis (GSEA) reveals the top 15 significantly dysregulated pathways in ARID2-deficient CRC cells. C) Gene Set Enrichment Analysis (GSEA) reveals a significant positive enrichment of oncogenic signalling pathways in HCT116 KO cells. Representative enrichment plots for the Wnt/β-catenin signalling, MYC targets, and TNF-A signalling hallmark gene sets are shown. D) Hierarchical clustering and heatmap representation of Hallmark genes identified via GSEA. The heatmap illustrates the distinct expression profiles of target genes significantly upregulated (red) or downregulated (blue) in KO compared to WT HCT116 cells.

---

**Table S1.**
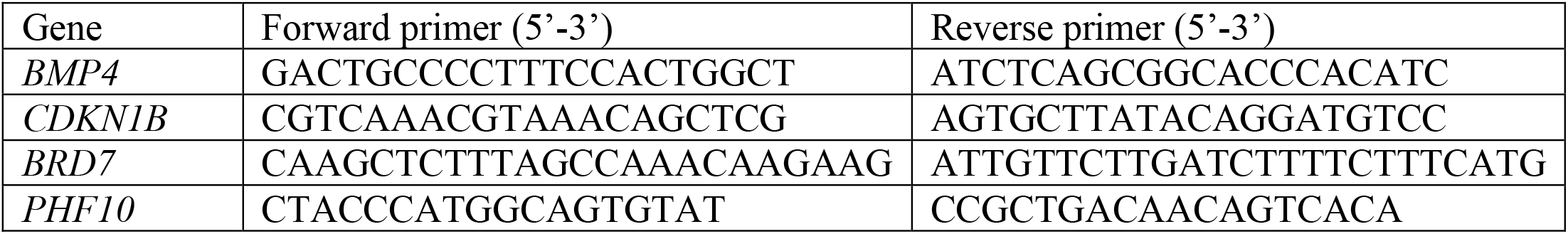
RT-qPCR primer sequences.

## References

1. Clapier CR, Iwasa J, Cairns BR, Peterson CL. Mechanisms of action and regulation of ATP-dependent chromatin-remodelling complexes. Nat Rev Mol Cell Biol. 2017 Jul;18(7):407–22. doi:10.1038/nrm.2017.26

2. Wilson BG, Roberts CWM. SWI/SNF nucleosome remodellers and cancer. Nat Rev Cancer. 2011 Jun 9;11(7):481–92. doi:10.1038/nrc3068 PubMed PMID: 21654818.

3. Gañez-Zapater A, Mackowiak SD, Guo Y, Tarbier M, Jordán-Pla A, Friedländer MR, et al. The SWI/SNF subunit BRG1 affects alternative splicing by changing RNA binding factor interactions with nascent RNA. Mol Genet Genomics MGG. 2022 Mar;297(2):463–84. doi:10.1007/s00438-022-01863-9 PubMed PMID: 35187582; PubMed Central PMCID: PMC8960663.

4. Kadoch C, Crabtree GR. Mammalian SWI/SNF chromatin remodeling complexes and cancer: Mechanistic insights gained from human genomics. Sci Adv. 2015 Jun 12;1(5):e1500447. doi:10.1126/sciadv.1500447

5. Chromatin remodelling during development | Nature [Internet]. [cited 2026 Mar 23]. Available from: https://www.nature.com/articles/nature08911

6. Loesch R, Chenane L, Colnot S. ARID2 Chromatin Remodeler in Hepatocellular Carcinoma. Cells. 2020 Sep 23;9(10):2152. doi:10.3390/cells9102152 PubMed PMID: 32977645; PubMed Central PMCID: PMC7598172.

7. Mashtalir N, D’Avino AR, Michel BC, Luo J, Pan J, Otto JE, et al. Modular Organization and Assembly of SWI/SNF Family Chromatin Remodeling Complexes. Cell. 2018 Nov 15;175(5):1272–1288.e20. doi:10.1016/j.cell.2018.09.032 PubMed PMID: 30343899; PubMed Central PMCID: PMC6791824.

8. The SWI/SNF complex in cancer — biology, biomarkers and therapy | Nature Reviews Clinical Oncology [Internet]. [cited 2026 Mar 24]. Available from: https://www.nature.com/articles/s41571-020-0357-3

9. Bala P, Singh AK, Kavadipula P, Kotapalli V, Sabarinathan R, Bashyam MD. Exome sequencing identifies ARID2 as a novel tumor suppressor in early-onset sporadic rectal cancer. Oncogene. 2021 Jan;40(4):863–74. doi:10.1038/s41388-020-01537-z PubMed PMID: 33262464.

10. Li M, Zhao H, Zhang X, Wood LD, Anders RA, Choti MA, et al. Inactivating mutations of the chromatin remodeling gene ARID2 in hepatocellular carcinoma. Nat Genet. 2011 Aug 7;43(9):828–9. doi:10.1038/ng.903 PubMed PMID: 21822264; PubMed Central PMCID: PMC3163746.

11. Duan Y, Tian L, Gao Q, Liang L, Zhang W, Yang Y, et al. Chromatin remodeling gene ARID2 targets cyclin D1 and cyclin E1 to suppress hepatoma cell progression. Oncotarget. 2016 Jul 19;7(29):45863–75. doi:10.18632/oncotarget.10244 PubMed PMID: 27351279; PubMed Central PMCID: PMC5216766.

12. Shi H, Tao T, Abraham BJ, Durbin AD, Zimmerman MW, Kadoch C, et al. ARID1A loss in neuroblastoma promotes the adrenergic-to-mesenchymal transition by regulating enhancer-mediated gene expression. Sci Adv. 2020 Jul 15;6(29):eaaz3440. doi:10.1126/sciadv.aaz3440

13. Meisenberg C, Pinder SI, Hopkins SR, Wooller SK, Benstead-Hume G, Pearl FMG, et al. Repression of Transcription at DNA Breaks Requires Cohesin throughout Interphase and Prevents Genome Instability. Mol Cell. 2019 Jan 17;73(2):212–223.e7. doi:10.1016/j.molcel.2018.11.001 PubMed PMID: 30554942; PubMed Central PMCID: PMC6344341.

14. de Castro RO, Previato L, Goitea V, Felberg A, Guiraldelli MF, Filiberti A, et al. The chromatin-remodeling subunit Baf200 promotes homology-directed DNA repair and regulates distinct chromatin-remodeling complexes. J Biol Chem. 2017 May 19;292(20):8459–71. doi:10.1074/jbc.M117.778183 PubMed PMID: 28381560; PubMed Central PMCID: PMC5437250.

15. An L, Jiang Y, Ng HHW, Man EPS, Chen J, Khoo US, et al. Dual-utility NLS drives RNF169-dependent DNA damage responses. Proc Natl Acad Sci. 2017 Apr 4;114(14):E2872–81. doi:10.1073/pnas.1616602114

16. Ran FA, Hsu PD, Wright J, Agarwala V, Scott DA, Zhang F. Genome engineering using the CRISPR-Cas9 system. Nat Protoc. 2013 Nov;8(11):2281–308. doi:10.1038/nprot.2013.143

17. Oba A, Shimada S, Akiyama Y, Nishikawaji T, Mogushi K, Ito H, et al. ARID2 modulates DNA damage response in human hepatocellular carcinoma cells. J Hepatol. 2017 May 1;66(5):942–51. doi:10.1016/j.jhep.2016.12.026

18. George SA, Kotapalli V, Ramaswamy P, Kumar R, Gowrishankar S, Uppin SG, et al. Novel oncogenic transcriptional targets of mutant p53 in esophageal squamous cell carcinoma. J Cell Biochem. 2024 Apr;125(4):e30534. doi:10.1002/jcb.30534 PubMed PMID: 38358025.

19. Animireddy S, Kavadipula P, Kotapalli V, Gowrishankar S, Rao S, Bashyam MD. Aberrant cytoplasmic localization of ARID1B activates ERK signaling and promotes oncogenesis. J Cell Sci. 2021 Feb 26;134(4):jcs251637. doi:10.1242/jcs.251637 PubMed PMID: 33443092.

20. Babraham Bioinformatics - FastQC A Quality Control tool for High Throughput Sequence Data [Internet]. [cited 2026 Mar 24]. Available from: https://www.bioinformatics.babraham.ac.uk/projects/fastqc/

21. Gupta A, Avadhanula S, Bashyam MD. Evaluation of the gene fusion landscape in early onset sporadic rectal cancer reveals association with chromatin architecture and genome stability. Oncogene. 2024 Aug;43(32):2449–62. doi:10.1038/s41388-024-03088-z PubMed PMID: 38937601.

22. Martin M. Cutadapt removes adapter sequences from high-throughput sequencing reads. EMBnet.journal. 2011 May 2;17(1):10–2. doi:10.14806/ej.17.1.200

23. Patro R, Duggal G, Kingsford C. Salmon: Accurate, Versatile and Ultrafast Quantification from RNA-seq Data using Lightweight-Alignment.

24. Soneson C, Love MI, Robinson MD. Differential analyses for RNA-seq: transcript-level estimates improve gene-level inferences. F1000Research. 2016 Feb 29;4:1521. doi:10.12688/f1000research.7563.2 PubMed PMID: 26925227; PubMed Central PMCID: PMC4712774.

25. Love MI, Huber W, Anders S. Moderated estimation of fold change and dispersion for RNA-seq data with DESeq2. Genome Biol. 2014;15(12):550. doi:10.1186/s13059-014-0550-8 PubMed PMID: 25516281; PubMed Central PMCID: PMC4302049.

26. Wu T, Hu E, Xu S, Chen M, Guo P, Dai Z, et al. clusterProfiler 4.0: A universal enrichment tool for interpreting omics data. Innov Camb Mass. 2021 Aug 28;2(3):100141. doi:10.1016/j.xinn.2021.100141 PubMed PMID: 34557778; PubMed Central PMCID: PMC8454663.

27. Huang DW, Sherman BT, Lempicki RA. Systematic and integrative analysis of large gene lists using DAVID bioinformatics resources. Nat Protoc. 2009;4(1):44–57. doi:10.1038/nprot.2008.211 PubMed PMID: 19131956.

28. Subramanian A, Tamayo P, Mootha VK, Mukherjee S, Ebert BL, Gillette MA, et al. Gene set enrichment analysis: A knowledge-based approach for interpreting genome-wide expression profiles. Proc Natl Acad Sci. 2005 Oct 25;102(43):15545–50. doi:10.1073/pnas.0506580102

29. Zhan T, Rindtorff N, Boutros M. Wnt signaling in cancer. Oncogene. 2017 Mar;36(11):1461–73. doi:10.1038/onc.2016.304

30. Das LP, Pitty RH, Asokan K, C L K, M S A, Ramanathan A. Analysis of ARID2 Gene Mutation in Oral Squamous Cell Carcinoma. Asian Pac J Cancer Prev APJCP. 2017 Oct 26;18(10):2679–81. doi:10.22034/APJCP.2017.18.10.2679 PubMed PMID: 29072391; PubMed Central PMCID: PMC5747389.

31. Moreno T, Monterde B, González-Silva L, Betancor-Fernández I, Revilla C, Agraz-Doblas A, et al. ARID2 deficiency promotes tumor progression and is associated with higher sensitivity to chemotherapy in lung cancer. Oncogene. 2021 Apr;40(16):2923– 35. doi:10.1038/s41388-021-01748-y PubMed PMID: 33742126; PubMed Central PMCID: PMC7610680.

32. Cajuso T, Hänninen UA, Kondelin J, Gylfe AE, Tanskanen T, Katainen R, et al. Exome sequencing reveals frequent inactivating mutations in ARID1A, ARID1B, ARID2 and ARID4A in microsatellite unstable colorectal cancer. Int J Cancer. 2014 Aug 1;135(3):611–23. doi:10.1002/ijc.28705 PubMed PMID: 24382590.

33. Carcamo S, Nguyen CB, Grossi E, Filipescu D, Alpsoy A, Dhiman A, et al. Altered BAF occupancy and transcription factor dynamics in PBAF-deficient melanoma. Cell Rep. 2022 Apr 5;39(1):110637. doi:10.1016/j.celrep.2022.110637 PubMed PMID: 35385731; PubMed Central PMCID: PMC9013128.

34. Shang L, Cho MT, Retterer K, Folk L, Humberson J, Rohena L, et al. Mutations in ARID2 are associated with intellectual disabilities. Neurogenetics. 2015 Oct;16(4):307–14. doi:10.1007/s10048-015-0454-0 PubMed PMID: 26238514.

35. Kang E, Kang M, Ju Y, Lee SJ, Lee YS, Woo DC, et al. Association between ARID2 and RAS-MAPK pathway in intellectual disability and short stature. J Med Genet. 2021 Nov;58(11):767–77. doi:10.1136/jmedgenet-2020-107111 PubMed PMID: 33051312.

36. Medina PP, Carretero J, Ballestar E, Angulo B, Lopez-Rios F, Esteller M, et al. Transcriptional targets of the chromatin-remodelling factor SMARCA4/BRG1 in lung cancer cells. Hum Mol Genet. 2005 Apr 1;14(7):973–82. doi:10.1093/hmg/ddi091 PubMed PMID: 15731117.

37. The SWI/SNF complex in cancer — biology, biomarkers and therapy | Nature Reviews Clinical Oncology [Internet]. [cited 2026 Mar 24]. Available from: https://www.nature.com/articles/s41571-020-0357-3

38. Dreier MR, Walia J, de la Serna IL. Targeting SWI/SNF Complexes in Cancer: Pharmacological Approaches and Implications. Epigenomes. 2024 Feb 4;8(1):7. doi:10.3390/epigenomes8010007 PubMed PMID: 38390898; PubMed Central PMCID: PMC10885108.

39. Hore P, Sarkar S, Bashyam MD. Evaluating the ‘ARID’ landscape of SWI/SNF complexes with relevance to cancer. Gene. 2025 Sep 15;965:149664. doi:10.1016/j.gene.2025.149664

